# Would an RRS by any other name sound as RAD?

**DOI:** 10.1101/283085

**Authors:** Erin O Campbell, Bryan M T Brunet, Julian R Dupuis, Felix A H Sperling

## Abstract

Sampling markers throughout a genome with restriction enzymes emerged in the 2000s as reduced representation shotgun sequencing (RRS). Rapid advances in sequencing technology have since spurred modifications of RRS, giving rise to many derivatives with unique names, such as RADseq. But naming conventions have often been more creative than consistent, with unclear criteria for recognition as a unique method resulting in a proliferation of names characterized by ambiguity. We conducted a literature review to assess methodological and etymological relationships among 36 restriction enzyme-based methods, as well as rates of correct referencing of commonly-used methods. We identify several instances of methodological convergence or misattribution in the literature, and note that many published derivatives have modified only minor elements of parent protocols. We urge greater restraint in naming derivative methods, to strike a better balance between clarity, recognition of scientific innovation, and correct attribution.

## INTRODUCTION

Recent advances in next-generation sequencing (NGS) have given researchers access to unprecedented amounts of genomic data. The versatility of NGS, exemplified by its myriad applications to biology (Andrews *et al.* 2016), is arguably one of its greatest assets and has in turn led to more than 400 published methods that use this technology (Hadfield & Retief 2018). While NGS has indisputably spurred rapid innovation across biology, associated names have also proliferated. These names are commonly acronyms meant to clearly identify a methodology or application but, due to their sheer numbers, are now themselves a source of confusion. A set of suggested guidelines for the use of such acronyms was published several years ago (NUAP 2011), but Hadfield & Retief (2018) have recently reignited this conversation, discussing the excess of names for NGS methods but not analyzing their patterns of naming, publication, or citation

Methods that use restriction enzymes to sample genomes represent an informative subset of NGS techniques to explore in this context. These methods provide diverse options for reducing genomic complexity and surveying large numbers of loci across populations or species, and are widely used in ecology and evolutionary biology (Baird *et al.* 2008; Davey *et al.* 2011). Early approaches include reduced representation shotgun sequencing (RRS, Altshuler *et al.* 2000) and Complexity Reduction of Polymorphic Sequences (CRoPs^™^, van Orsouw *et al.* 2007), which have since served as springboards for derivative techniques, most of them published with unique names. At least 36 of these methods have been published as of December 2017 (Table S1).

Two methods in particular, Restriction site-associated DNA sequencing (RADseq, Baird *et al.* 2008) and Genotyping-by-Sequencing (GBS, Elshire *et al.* 2011), have been modified for diverse work on association and genetic mapping, population structure, and shallow-scale phylogenetic relationships (Poland *et al.*, 2012; Baird *et al.*, 2013; Narum *et al.*, 2013; Eaton, 2014). They are so popular that 26 methods have explicitly been modified from either RADseq or GBS (Table S1), with their increasing importance demonstrated by recent reviews (Davey *et al.*, 2011; Andrews *et al*., 2016; Jiang *et al.* 2016), as well as debates (e.g. Andrews & Luikart (2014) and Puritz *et al.* (2014), and Lowry *et al.* (2016, 2017), McKinney *et al.* (2017), and Catchen *et al.* (2017)). Although reviews have tried to distinguish these approaches, it is clear from these publications as well as informal discussion on online forums (Table S2) that differences between many techniques are perceived to be minor and subtle. Naming conventions for derivatives have been variable and inconsistent, and literature discussing or employing these techniques has been ambiguous about the origins of techniques as well as which names to use as “catch-all” terms.

For instance, “GBS” is sometimes used to refer to all restriction-based methods collectively (e.g. Franchini *et al.* 2017), while other authors take the opposite approach and use “RADseq” as the generic term (e.g. Hoffberg *et al.* 2016). Two-enzyme, or double digest, adaptations of these techniques are similarly ambiguous; Peterson *et al.* (2012) has been credited with developing this approach (see Andrews *et al.* 2016) and did coin the term ddRAD to expand on the already-popular single-enzyme RADseq approach (Baird *et al*. 2008), but the CRoPS^™^ protocol (van Orsouw *et al.* 2007) was the first RRS method to do this. Some authors even give a common acronym a new meaning. For example, NextRAD (Fu *et al.* 2017) was developed by authors of the original RADseq papers (Miller *et al.* 2007; Baird *et al.* 2008) but uses RAD in this derived protocol to stand for Reductively-Amplified DNA. Thus, reduced representation genome sampling methods (hereafter referred to as RRS methods after Altshuler *et al.* 2000) exemplify the naming problem that is now typical of NGS methods.

Given the proliferation of RRS methods and ambiguity of their naming conventions (Jiang *et al.* 2016), we have sought to characterize RRS methods in a literature review and meta-analysis. We asked two main questions. First, what are the trends or criteria for naming new methods? And second, are researchers citing and referring to methods correctly? To answer these questions, we summarized the methodological and etymological relationships of these techniques in a concept map, and then conducted a meta-analysis of citation metrics to investigate patterns of literature referencing.

## METHODS

### Literature review and concept mapping

We compiled a list of RRS methods published on or before 31 December 2017 (N = 36), and evaluated approaches based on their methodological characteristics (Table S2). We then created a conceptual map of all methods, linking each derived technique to the main protocol that served as the basis for the modification, as specified by the authors (Fig. 1). In several cases a parent protocol was not directly specified, and in these instances, we linked methods based on overall methodological similarity. The subjective construction of this map reflects our experience as typical arms-length users of several of these approaches. Any technique that explicitly altered a protocol was considered a direct modification, and in this conceptualized map, a separate node. We plotted defining characteristics for each derivative along the connecting branches to assess distinctiveness or methodological convergence. Defining characteristics were generally those considered by the authors of the protocol to distinguish the derived method from its parent. To preserve clarity, characteristics that were highly variable across methods (for instance barcode and adaptor design and the overall order of methodological steps in each protocol) were not plotted on the map unless they were definitive for the method(s). We also downloaded complete citation data from Web of Science® for the 36 methods, and determined the average number of citations per year for each publication. The size of ellipses in Fig. 1 reflects these numbers.

**Figure 1.**
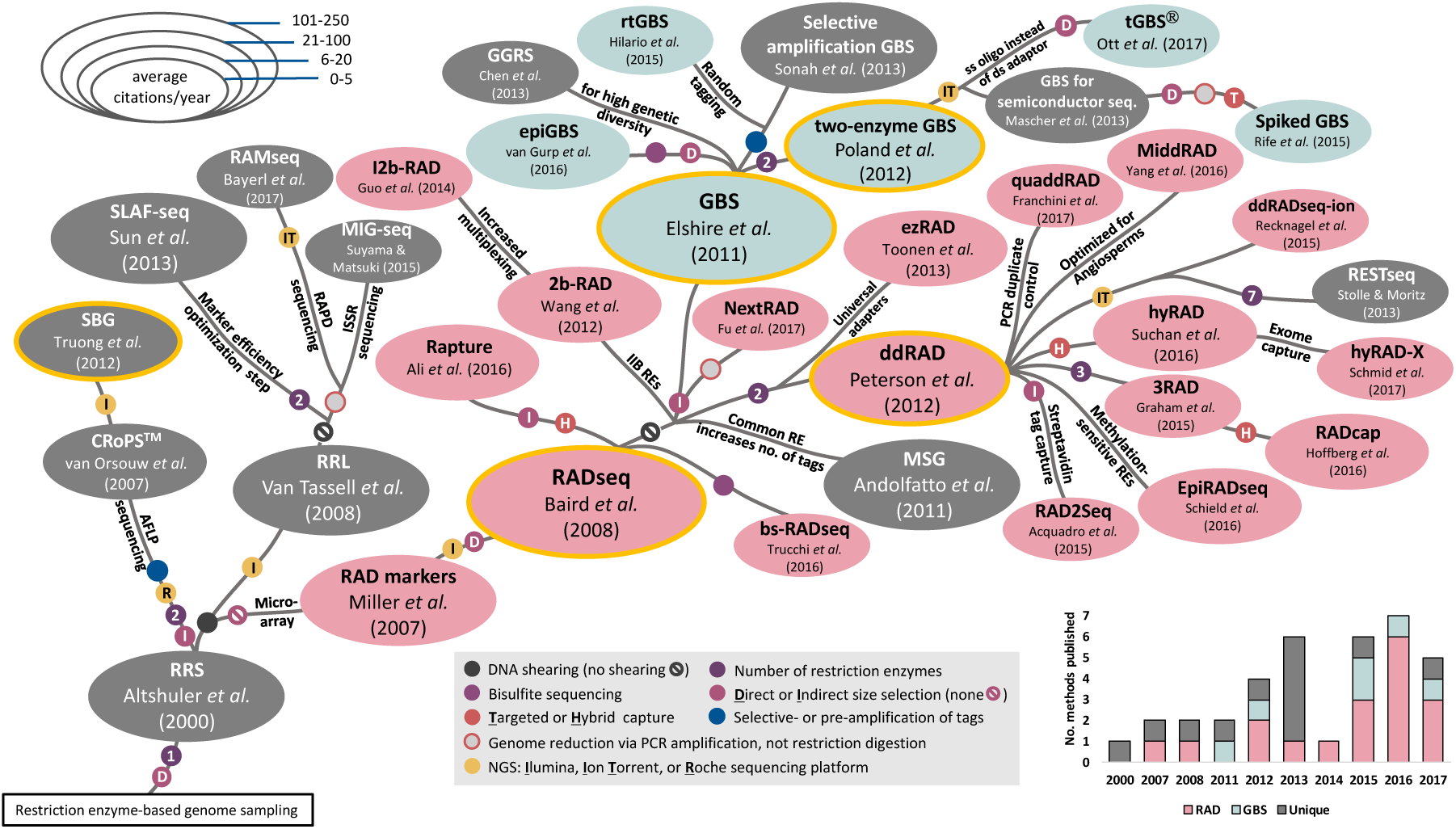
Concept map displaying methodological and etymological relationships among 36 reduced representation genome-sampling methods and their derivatives. Branches connect derived methods to their inferred parent protocols, and significant differences between protocols are indicated by coloured circles on branches. Variations that originate only once are indicated by text along branches. Red ellipses indicate named methods using the “RAD” acronym, blue ellipses indicate names derived from “GBS”, and methods with unique names, or lacking names altogether, are in grey. The five methods used to assess attribution rates in an accompanying literature review (Fig. 2) are indicated by ellipses with a gold outline. Inset histogram shows accumulation of methods by year. See Table S1 for a summary of each method.

### Meta-analysis of commonly used RRS approaches

To assess whether methods are recognized and attributed accurately in the literature, we reviewed all journal articles citing one- and two-enzyme RADseq and GBS (RADseq, Baird *et al.* 2008; GBS, Elshire *et al.* 2011; two-enzyme GBS, Poland *et al.* 2012; and ddRAD, Peterson *et al.* 2012), as well as Sequence-based Genotyping (SBG, Truong *et al.* 2012). RADseq, GBS, two-enzyme GBS and ddRAD are four of the most widely cited RRS approaches, and have been extensively modified to form the basis of many derivative methods. While the SBG protocol of Truong *et al.* (2012) is far less frequently cited, we included it for its methodological and etymological similarity to these methods as well as its date of publication, which occurred between that of Poland *et al.* (2012) and Peterson *et al.* (2012). It is also the subject of U.S. patent 8,815,512 B2, owned by KeyGene, which claims legal ownership and protection of all methods that simultaneously discover and genotype single nucleotide polymorphisms, including RADseq, GBS, two-enzyme GBS and ddRAD (KeyGene 2016).

Complete citation lists were downloaded from Web of Science® on 6 February 2018 for the period up to and including 31 December 2017 for each of Baird *et al.* (2008), Elshire *et al.* (2011), Poland *et al.* (2012), Truong *et al.* (2012), and Peterson *et al.* (2012). The lists were then combined and filtered to remove duplicates (Table S3). Only articles whose titles, abstracts, or keywords contained “GBS”, “SBG” or “RAD” (and all variant search strings in Table S4) were retained for further analysis. Incorrect name usage was defined as any case of an alternate name being used to refer to a technique (e.g. “RAD” to describe the GBS protocol of Elshire *et al.* (2011)). Strings for “two-enzyme GBS” and “ddRAD” were not searched separately since these were treated as variants of “GBS” and “RAD”, respectively. A complete description of the methods used in the literature review is in Table S6.

## RESULTS

### Construction of RRS conceptual map

Of the 36 RRS methods we examined, those of Baird *et al.* (2008), Peterson *et al.* (2012), and Elshire *et al.* (2011) are the precursors of the greatest number of directly derived methods (Fig. 1), and the most highly cited (Table S2). “RAD” was used in 18 named techniques, while “GBS” was used in 6 and the remaining 12 methods had names that lacked “RAD”, “GBS”, or any specified name at all (see Sonah *et al.* 2013 and Mascher *et al.* 2013). Many derived methods were named after the protocol they modified (*e.g.*: ddRAD (Peterson *et al.* 2012) from RADseq (Baird *et al.* 2008)), but there were several exceptions (*e.g.*, SBG (Truong *et al.* 2012) from CRoPS^™^ (van Orsouw *et al.* 2007)). We observed multiple occurrences of methodological convergence across methods, including the use of paired restriction enzymes in double digest methods, sequence capture, bisulfite sequencing, and the use of PCR amplification to create reduced representation libraries, which we discuss below.

### Meta-analysis of GBS, SBG and RAD citation accuracy

The number of journal articles that refer to “GBS”, “SBG”, or “RAD” within their title, abstract or keywords and uniquely cite either Baird *et al.* (2008), Elshire *et al.* (2011), Poland *et al.* (2012), Truong *et al.* (2012), or Peterson *et al.* (2012) has increased rapidly since 2010, with the greatest number of citations occurring in 2017. Of a total of 788 journal articles, 335 (42.5%) refer only to GBS, 2 (0.2%) refer only to SBG, and 418 (53.1%) refer only to RAD (Fig. 2; Table S5). Two or more of these names (“Multiple(≥2)”) are used in only 33 (4.3%) journal articles and these refer only to GBS and RAD, not SBG (Fig. 2; Table S5).

**Figure 2.**
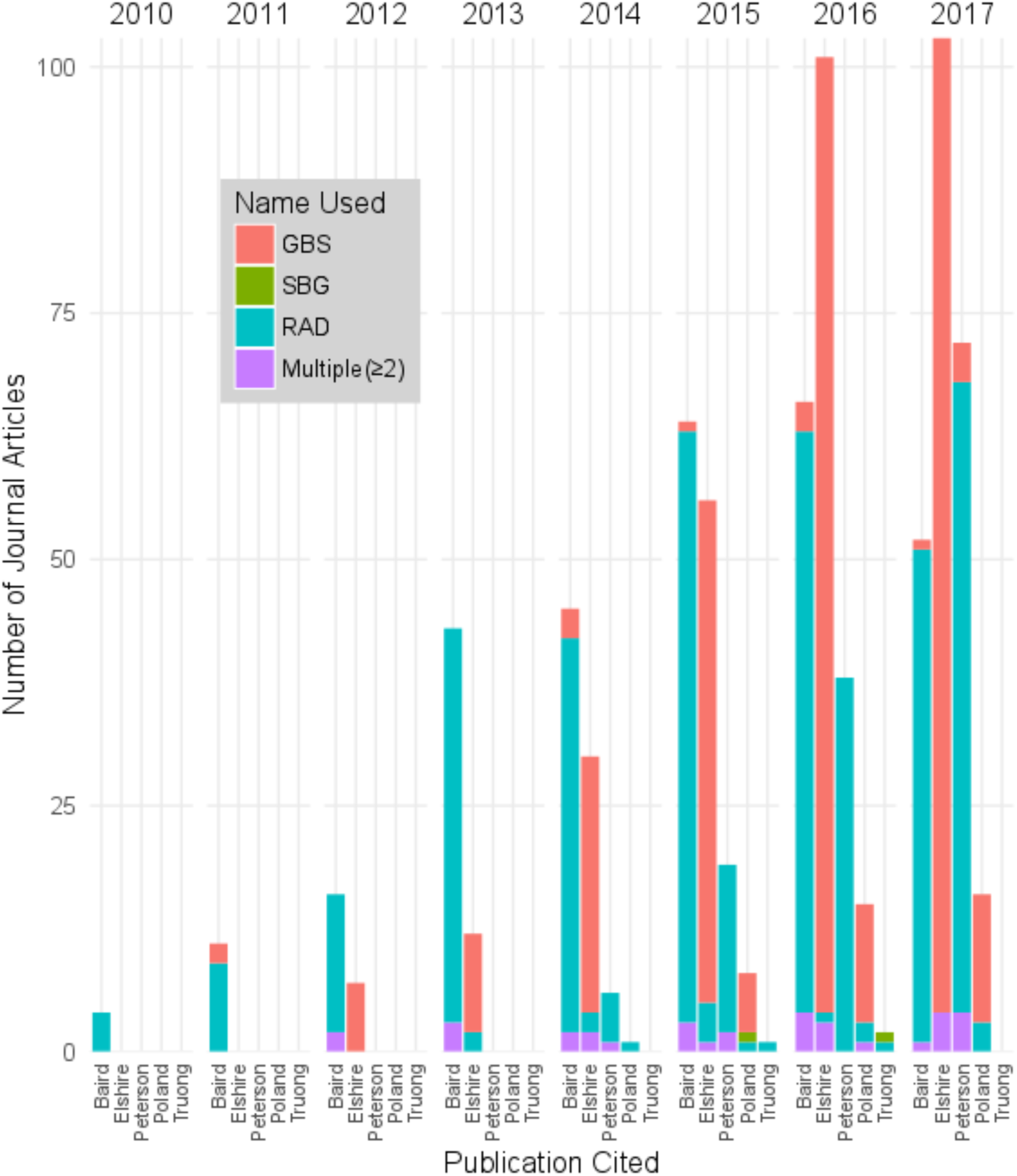
Trends in the use of the names “GBS” (from Elshire *et al.* (2011) and Poland *et al.* (2012)), “SBG” (from Truong *et al*. (2012)), and “RAD” (from Baird *et al.* (2008) and Peterson *et al.* (2012)) in the title, abstract, or keywords of journal articles that cite either Baird *et al.* (2008), Elshire *et al.* (2011), Poland *et al.* (2012), Truong *et al.* (2012) or Peterson *et al.* (2012). Bars indicate the number of journal articles citing each publication, while colours indicate the number referring to each name. About 8.4% of papers use an ambiguous or incorrect name in reference to the cited method (e.g. ∼4% of papers uniquely citing Baird *et al*. (2008) in 2017 refer specifically to GBS or SBG alone or in combination with RAD, despite neither name being used in that paper).

Each name has been used inconsistently to refer to methods described by their parent publications, but to varying degrees: 8% (28/349) of publications that uniquely cite Elshire *et al.* (2011) or Poland *et al.* (2012) refer to SBG or RAD alone or in combination with GBS; 66.7% (2/3) of publications that uniquely cite Truong *et al.* (2012) refer to GBS or RAD alone or in combination with SBG; and 8.3% (36/436) of publications that uniquely cite Baird *et al.* (2008) or Peterson *et al.* (2012) refer to GBS or SBG alone or in combination with RAD (Fig. 2; Tables S3 and S5). Thus, use of ambiguous or incorrect names are apparent in about 8.4% of journal articles citing these five papers.

## DISCUSSION

We have applied a review and literature meta-analysis to characterize the naming and use of RRS methods. Our concept map shows that RAD-based methods are more numerous than GBS-based methods. Although derived methods are often given unique names, most follow some of the etymological elements of the parent technique that was modified, even when derived protocols from different camps converge methodologically (Fig. 1). We also identified a rate of ∼8.4% ambiguous or incorrect citations for these methods (Fig. 2).

The RAD acronym leads the popularity race when considering citations for RAD-based methods as well as the number of derivative protocols bearing this term; GBS-based methods have fewer overall citations and methodological offspring. While the original RAD or modified ddRAD methods may simply be more methodologically attractive, unconscious linguistic factors in the naming of derivatives may also be contributing to this trend. Acronyms that form simple, recognizable words are more likely to be remembered (NUAP 2011), so this may explain use of the RAD acronym despite citation of a GBS or SBG publication (Fig. 2). RAD has also proven to be easy to incorporate into memorable titles that improve name recognition and visibility in a rapidly expanding field (*e.g.* “Demystifying the RAD fad” (Puritz *et al.* 2014); “Breaking RAD” (Lowry *et al.* 2016); present study). However, rates of potential misattribution do not appear to be biased toward RAD over GBS (Fig. 2), and so researchers who are unclear or unconvinced of the distinctions between methods may simply be randomly using both terms as synonyms. Methodological convergence by several GBS- and RAD-based techniques (Fig. 1) could contribute to further ambiguity among methods.

*“What’s in a name?”* (Shakespeare 1594-98). Separate publication of a method implies that the authors consider the new method to be substantively different from other published methods, thereby warranting a separate name. But for RRS methods, differences between many techniques are minor, often primarily implementing streamlined library preparation and cost reduction (e.g.: GGRS (Chen *et al.* 2013), ezRAD (Toonen *et al.* 2013)), or adaptations that optimize methods for specific groups of organisms (e.g.: MiddRAD, Yang *et al.* 2016). Many published methods arguably do not meet this criterion for publication (NUAP 2011). And while some RRS methods have made larger methodological changes, for instance the use of sequence capture or bisulfite sequencing, the publication of these methods is problematic for other reasons.

Naming of a method also suggests ownership over that method (NUAP 2011). In cases where only minor changes were made to an existing protocol, the authors of the new method profit from advances made by prior authors, which may comprise the bulk of the methodology. The broad convergence of several methods in Fig. 1 creates an additional layer of complexity, as two separate groups of authors are essentially claiming ownership over similar techniques that have different names. For instance, ddRADseq-ion (Recknagel *et al.* 2015) and GBS for semiconductor sequencing platforms (Mascher *et al.* 2013) have both incorporated double digests and modified adaptors for Ion Torrent sequencing. EpiGBS (van Gurp *et al.* 2016) and bsRADseq (Trucchi *et al.* 2016) both incorporate bisulfite sequencing, and several methods have employed some form of sequence capture (Spiked GBS (Rife *et al.* 2016); RADcap (Hoffberg *et al.* 2016); HyRAD (Suchan *et al.* 2016); Rapture (Ali *et al.* 2016); 3RAD (Graham *et al.* 2015); hyRADx (Schmid *et al.* 2017)).

While each technique is prone to distinct biases and technical difficulties (van Dijk *et al.* 2014; Flanagan & Jones 2017), we argue that most RRS methods are sufficiently similar that they can be adapted to a user-specified combination of characteristics that suit the needs of an individual study. The recently upheld US KeyGene patent covering these methods also seems to suggest that, at least from a legal standpoint, they are not significantly different from one another (US Patent 8,815,512 B2). So why do we keep naming derivative methods, and how can we work to preserve clarity of communication in discussing them?

*“Action is eloquence”* (Shakespeare 1605-08). Rapid sequencing advances may have unwittingly created a sense of momentum among researchers, thereby fostering the proliferation of names for genome-sampling techniques. Almost half of the methods in Fig. 1 and all five of the key methods in Fig. 2 were published in *PLoS ONE*, which has published several RRS derivatives within the same years. Other journals have also published multiple derivative methods, although to a lesser degree. Several researchers have also been involved the naming and publication of more than one method, indicating research groups developing suites of techniques.

This suggests a preoccupation with name recognition as a means to increase the visibility of research. We recognize that catchy titles are not inherently negative or irresponsible, and that this practice can beneficially increase the impact of research. But the recent “modify, name, and publish” trend seems more likely to be driven by efforts to increase citations, which dilutes the eloquence of acronyms. This system further confers risk to researchers who choose not to name an adapted technique, by leaving a door open for someone else to employ the same change and name it, taking the credit. Because academic success is so closely tied to citation metrics, there is little incentive to take the high road.

It is instructive to compare reliance on easy-to-digest acronyms to online clickbait headlines in academic publishing and research. An example may be the recent publication of an incendiary essay presumably to increase the impact of a journal despite the article not passing peer review (Flaherty 2017). Academic metrics do not distinguish between “good” and “bad” citations (Gallien & Roelofs 2017), and we are incentivized to market our research beyond the merit of the research itself. Sequencing technology will undoubtedly continue to advance (Goodwin *et al.* 2016), and new RRS approaches will continue to evolve. Exploring the utility and limitations of these approaches has resulted in a wealth of biological knowledge that has been hitherto out of reach. At this level, Shakespeare’s immortal phrase got it right: “a rose by any other name would smell as sweet” (Shakespeare 1591-94). But Linnaeus may have disagreed with this sentiment – names *do* matter because they serve to communicate and organize the world around us.

We add our voices to those of Hadfield & Retief (2018); our scientific community would be better served by greater restraint in naming new techniques, except for indisputably large methodological innovations. Continued adaptation of methods is clearly beneficial, but the publication of new names for minor changes in existing NGS methodologies is a symptom of a larger cultural shift in academia. And the responsibility for righting that course lies with us as researchers, editors, and publishers.

## AUTHOR CONTRIBUTIONS

EOC, BMTB, JRD, and FAHS contributed to the study design and draft revisions. EOC and BMTB conducted the literature review and meta-analysis, and wrote the manuscript.

## ACKNOWLEDGEMENTS

This work was supported in part by an NSERC Discovery Grant to FAH Sperling (RES0016259).

